# Mechanical efficiency during sub-maximal cycling is underestimated because negative muscular power is ignored

**DOI:** 10.1101/2025.01.21.634028

**Authors:** Roeland T. Vos, Maarten F. Bobbert, Dinant A. Kistemaker

## Abstract

The *in vivo* mechanical efficiency of muscles has often been estimated during sub-maximal cycling. In this approach, it has implicitly been assumed that the average amount of positive mechanical muscle power equals the average mechanical power output, i.e., that no power is dissipated by muscles. Here, we investigated the validity of this assumption using an optimal control musculoskeletal model. We identified optimal muscle stimulation patterns for 4 cadences (60, 80, 100 and 120RPM) and 5 levels of average mechanical power output (50, 100, 150, 200 and 250W). We found that the amount of negative mechanical muscular power was substantial, with the average across all conditions being -84,6W (56,4%). The amount of negative mechanical muscular power was found to increase with increasing cadence and was independent of the average mechanical power output. To investigate the effect of negative muscular power on *in vivo* estimates of the muscular efficiency, we used our simulation results to correct gross efficiencies measured during sub-maximal cycling. The resulting increases in the gross efficiency were substantial, with the average increasing from 16.9% to 27.5%. These results suggest that current estimates of the muscular efficiency during sub-maximal cycling underestimate the true muscular efficiency.

## 1 Introduction

In the literature, estimates have been reported for the mechanical efficiency of human muscles during *in vivo* tasks. The muscular mechanical efficiency is defined as the ratio between the mechanical power produced by the muscles and the rate of metabolic energy consumed by the muscles to produce mechanical power (Ettema & Lorås, 2009; van Ingen Schenau et al., 1997). In general, the *in vivo* muscular mechanical efficiency cannot be measured directly, as it is not possible to measure the mechanical power and metabolic energy rate of the individual muscles. Therefore, estimates of the *in vivo* muscular mechanical efficiency commonly used measurements during sub-maximal tasks of whole-body oxygen consumption (V^̇^O_2_), carbon dioxide production (V^̇^CO_2_) and the externally delivered mechanical power (Ettema & Lorås, 2009; van Ingen Schenau et al., 1997).

Cycling is frequently used as a task in studies on *in vivo* muscular mechanical efficiency, because the externally delivered mechanical power can readily be determined by multiplying the torque delivered to the crank by the participant and the crank angular velocity. In many of these studies, values for the gross efficiency (GE) were reported (Chavarren & Calbet, 1999; Moseley et al., 2004; Samozino et al., 2006). GE by definition underestimates the true muscular mechanical efficiency. However, base-line subtractions used to ‘correct’ the metabolic energy rate, such as those used in the definitions of net efficiency or work efficiency, often result in unreasonably high mechanical efficiencies (Stainbsy et al., 1980). As summarized by Ettema and Lorås (2009) in their review, GE during sub-maximal cycling appears to depend mainly on two factors, namely the cadence and the average mechanical power output (AMPO) of the participant. For a given AMPO, increasing the cadence tends to lead to a decrease in GE (Chavarren & Calbet, 1999; Gaesser & Brooks, 1975; Sallet et al., 2006; Samozino et al., 2006; Sidossis et al., 1992), whereas increasing AMPO for a given cadence results in an increase in GE (Chavarren & Calbet, 1999; Moseley & Jeukendrup, 2001; Moseley et al., 2004; Samozino et al., 2006). The effects of cadence and AMPO on GE are illustrated in Fig. 1.

**Figure 1.**
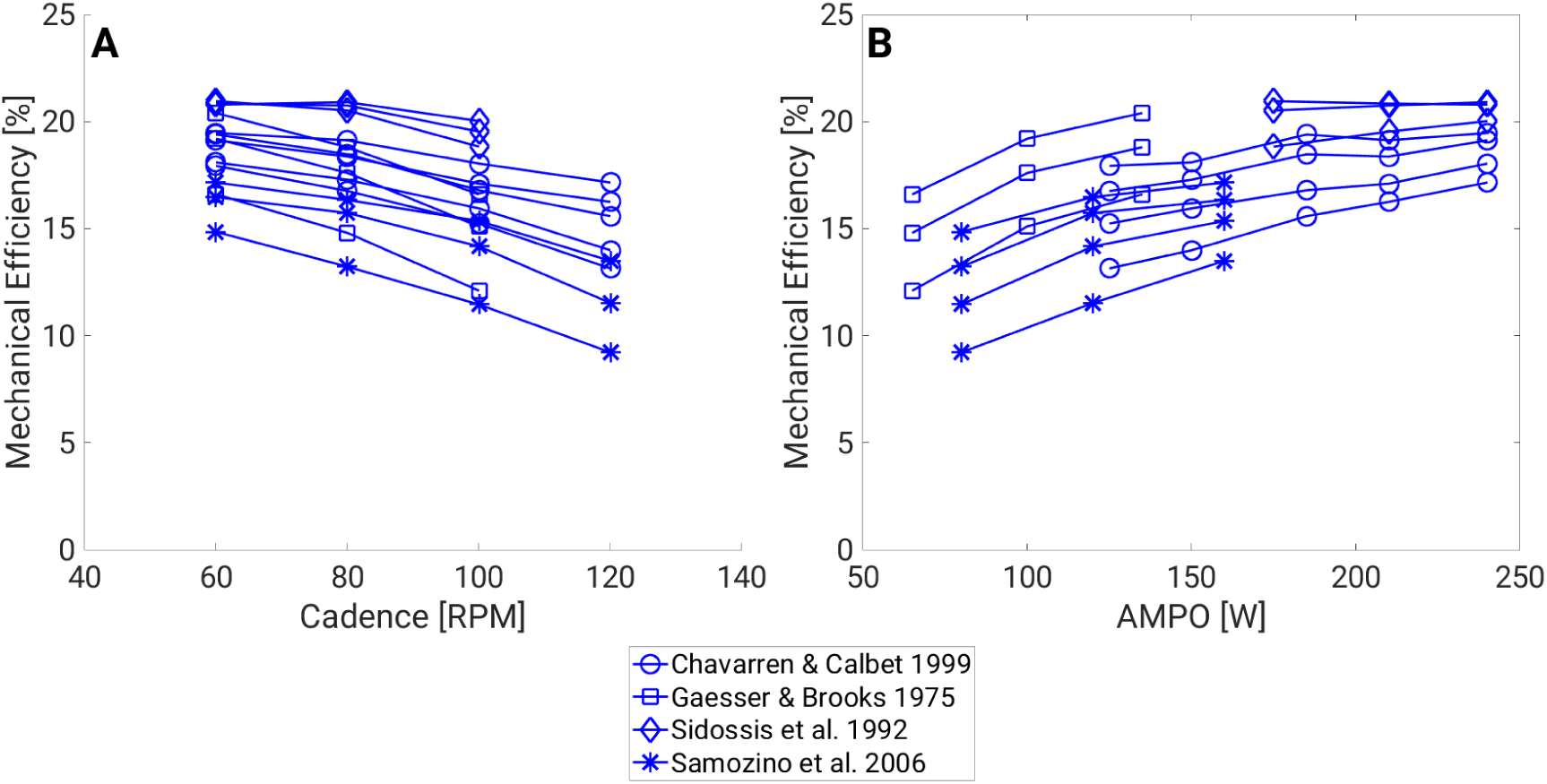
GE as a Function of Cadence and AMPO. GE obtained during sub-maximal-cycling cycling, plotted as a function of the cadence (**A**) or AMPO (**B**). Represented studies are Chavarren and Calbet (1999), Gaesser and Brooks (1975), Samozino et al. (2006), and Sidossis et al. (1992).

An implicit assumption made in the calculation of mechanical efficiency during sub-maximal cycling is that the muscles produce no negative mechanical power. This assumption is important, because only then does AMPO equal the average amount of positive muscular mechanical power (AMPO^+^). Results from simulation studies on both sprint and sub-maximal cycling suggest that the muscles produce non-negligible amounts of negative mechanical power (Kistemaker et al., 2023; Neptune & van den Bogert, 1997; van Soest & Casius, 2000). For example, Kistemaker et al. (2023) estimated that the power dissipation ratio (PDR) was about -0.20 in sprint cycling, which means that for every watt of positive mechanical muscular power the muscles dissipate 0.20 watt of negative power. To attain a desired AMPO, additional positive mechanical muscular power would be needed to compensate for negative mechanical muscular power. This additional positive mechanical power would lead to an increase in the metabolic power consumption. The muscles would also consume metabolic power for the production of negative mechanical power. If the assumption of no negative mechanical power were false, then the true muscular mechanical efficiency would be underestimated due to an underestimation of AMPO^+^ and an overestimation of the metabolic power consumed by the muscles. As far as we are aware, no studies have corrected their estimates of *in vivo* muscular mechanical efficiency during sub-maximal cycling for the effects of negative mechanical muscular power.

There is evidence that the average amount of negative mechanical muscular power (AMPO^−^) increases with increasing cadence due to the fact that the muscle activation dynamics are relatively slow (Neptune & Herzog, 2000; van Soest & Casius, 2000). As explained by van Soest and Casius (2000), in sprint cycling, attaining a high level of muscle active state sufficiently early in the shortening phase of the muscle requires that the muscle be stimulated in an earlier part of the cycle. This could lead to muscle stimulation starting during the end of the preceding lengthening phase and hence to the production of negative mechanical muscular power.

Imposed AMPO may also influence AMPO^−^. The increase in GE with increasing AMPO has primarily been attributed to the diminishing influence of the resting metabolic energy rate (Ettema & Lorås, 2009; Powers et al., 1984; Samozino et al., 2006). Currently, no studies have investigated a potential relationship between imposed AMPO and AMPO^−^ during sub-maximal cycling. Based on properties of the muscle activation dynamics, we would expect that an increase in AMPO would lead to an increase in AMPO^−^.

To obtain estimates of AMPO^−^ during sub-maximal cycling, one can use simulations of a musculoskeletal model of a cyclist, as the muscle force and contraction velocity can readily be obtained from the results of these simulations. Simulations with such models typically require optimizations of the input variables, i.e. muscle stimulation over time (STIM(*t*)), for a particular cost function. In contrast to sprint cycling, where the objective is unambiguous (maximizing AMPO), it is not clear on the basis of what criterion humans choose their muscle stimulation patterns during sub-maximal cycling. For example, for sub-maximal cycling it would seem logical to maximize mechanical efficiency. However, this contradicts the experimental finding that cyclists tend to prefer cadences higher than those that are optimal for efficiency (Hagberg et al., 1981; Marsh & Martin, 1993). Gidley et al. (2019) have compared a number of cost functions for simulating sub-maximal cycling and found that a cost function related to muscle fatigue (Ackermann & van den Bogert, 2010) led to the best overall match of simulation results with experimental data of human sub-maximal cycling.

The aims in the present study were as follows: first, we aimed to estimate AMPO^−^ during sub-maximal cycling at different combinations of cadence and AMPO. To do so, we used a simulation model that was able to capture the important characteristics of human sprint cycling (Bobbert et al., 2020; Kistemaker et al., 2023; van Soest & Casius, 2000) and optimal control techniques (see Kistemaker et al. (2023)) to identify optimal STIM(*t*), using a cost function based on muscle fatigue (Gidley et al., 2019). Using the results from these simulations, we also aimed to estimate the influence of AMPO^−^ on *in vivo* estimates of mechanical efficiency and re-examine the relationship between cadence, AMPO and GE in sub-maximal cycling.

## 2 Methods

### 2.1 Overview

Our first aim was to quantify AMPO^−^ during sub-maximal cycling. Using an optimal control musculoskeletal model of a human cyclist (Kistemaker et al., 2023), we simulated sub-maximal cycling at different combinations of AMPO and cadence. 4 cadences were used, ranging from 60 to 120 revolutions per minute, and 5 values for AMPO, which varied from 50 to 250 watts. The inputs of the model, STIM(*t*), were numerically optimized with respect to a cost function related to muscle fatigue (Ackermann & van den Bogert, 2010; Gidley et al., 2019). From these simulation results, we calculated AMPO^−^ and AMPO^+^ over one full cycle for each condition. In order to quantify the influence of negative mechanical muscular power on *in vivo* estimates of muscular mechanical efficiency, we re-estimated the experimentally observed GE in studies on sub-maximal cycling using AMPO^−^ and AMPO^+^.

### 2.2 Musculoskeletal model

The model, shown in Fig. 2, was two-dimensional and consisted of 5 segments: crank, foot, shank, thigh and a HAT-segment representing the head, arms and trunk combined. The model was driven by nine Hill-type muscle models. Parameter values were the same as those used by Kistemaker et al. (2023). The crank angular velocity was held constant throughout the cycle, as was done in previous studies. This meant that only one leg needed to be simulated as the legs are mechanically decoupled, thus simplifying the numerical optimization procedure as explained in Kistemaker et al. (2023).

**Figure 2.**
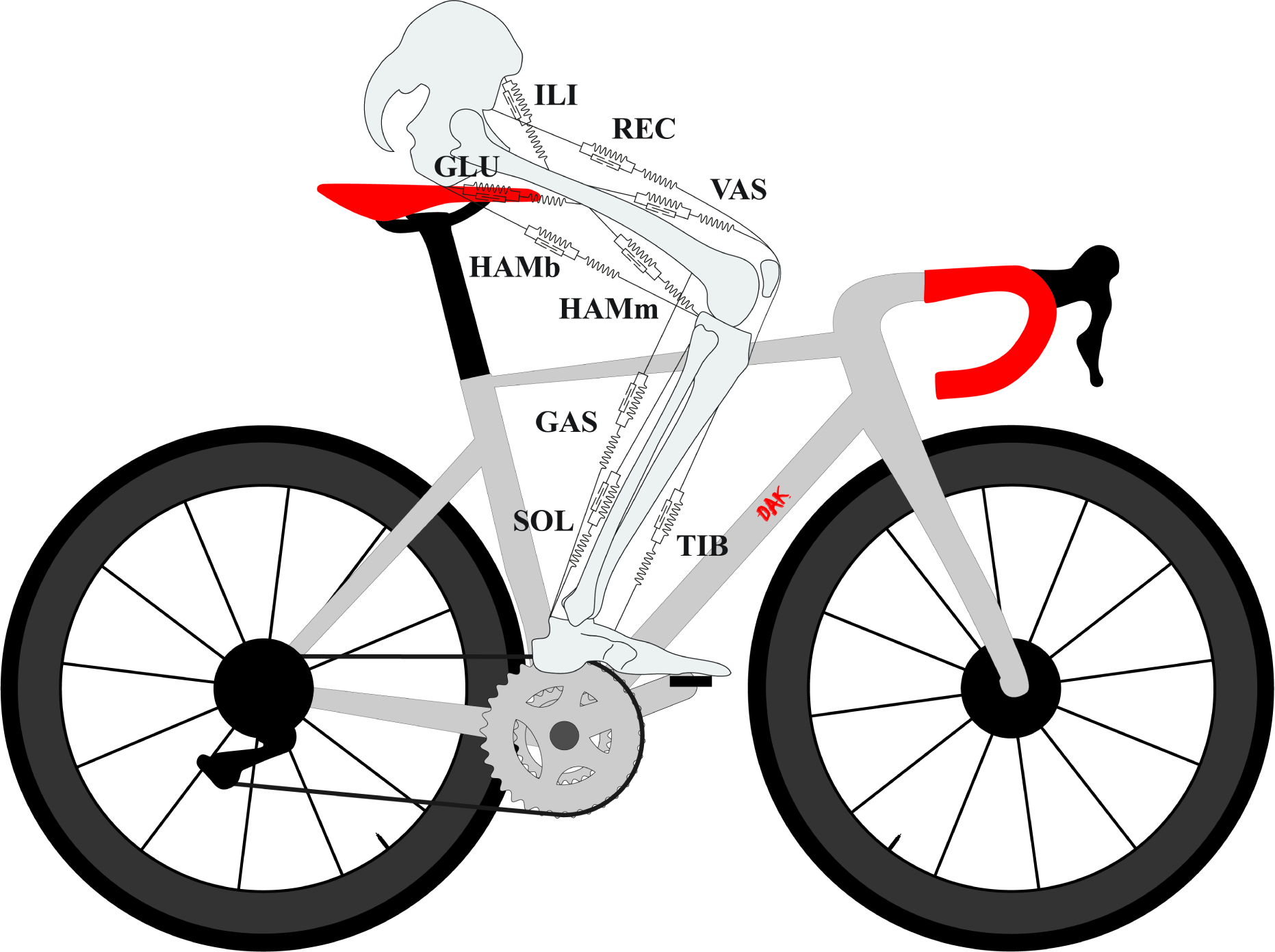
Musculoskeletal Model of a Human Cyclist.

### 2.3 Numerical optimization procedure

Optimal STIM(*t*) for one full cycle were found through the use of the sparse nonlinear optimal controller (SNOPT; TOMLAB Optimization, Pullman, WA). Model inputs and state variables were discretized and model dynamics were expressed as constraint equations in a so-called direct collocation approach (Betts, 2010). Periodicity constraints were added on the states of the musculoskeletal model to ensure that the initial and final values of the states were equal. Furthermore, the cadence and AMPO were imposed through additional constraints. Similar to Kistemaker et al. (2023), 10 optimization rounds per condition were performed. Each round was started from a random initial guess, and the results from the round with the best value for the cost function were used for further analysis.

The cost function used for the optimizations was based on muscle fatigue (Ackermann & van den Bogert, 2010; Gidley et al., 2019). The cost function was defined as follows:

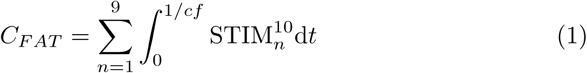

Where *cf* was the cycling frequency in hertz and *n* the number of muscles.

### 2.4 AMPO^+^ and AMPO*^−^*

For each condition we calculated AMPO^+^ and AMPO^−^ over one full cycle. AMPO^+^ and AMPO^−^ were defined as follows:

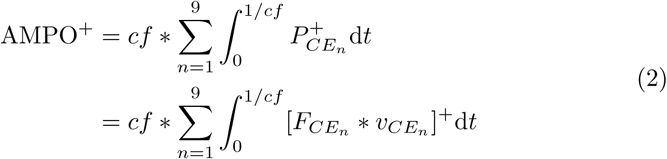

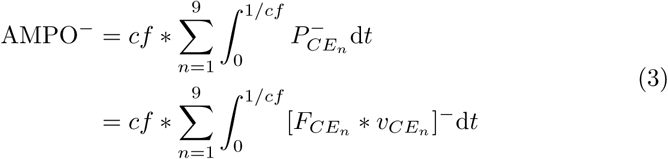

Where 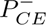 was the instantaneous negative mechanical power of the contractile element and *F_CE_* and *v_CE_* the force and contraction velocity of the contractile element.

### 2.5 Effect of AMPO*^−^* on *in vivo* mechanical efficiency estimates

Using the results from our simulations, we have re-estimated GE values obtained during sub-maximal cycling by accounting for the effects of AMPO^−^. The first correction was to subtract the metabolic power consumed for the production of negative mechanical muscular power from the experimentally measured metabolic power. For each GE obtained from the literature, we calculated AMPO^−^ for the corresponding combination of cadence and AMPO using our simulation results. Repeating the numerical optimizations for each of these combinations of cadence and AMPO would lead to an inconsistent set of combinations, because not all used cadences would be combined with all values of AMPO, and vice versa. To avoid this, we applied linear interpolation to the AMPO^−^ values obtained from the set of combinations described in the Overview. The metabolic power used for AMPO^−^ was then estimated by using the observation that muscles have a mechanical efficiency of -120% while producing negative mechanical power (Abbott et al., 1952). GE could then be re-calculated using the corrected metabolic power estimate. The second step was then to take the corrected GE from the first step and multiply it with the ratio between AMPO^+^ and AMPO. This was done to correct for additional positive mechanical muscular power due to negative mechanical muscular power.

## 3 Results

### 3.1 Optimization runs converged to similar solutions

For all simulated conditions, we were able to find solutions. To check for the convergence of the optimizations, we first summed STIM(*t*) per condition for both the best and second best solution. We then calculated the root mean square difference between the best and second best solutions in terms of the summed stimulations. The root mean square difference was only 2.8, indicating that the optimizations converged successfully.

### 3.2 AMPO*^−^* was substantial

Fig. 3 shows AMPO^+^ and AMPO^−^, for a subset of the conditions and plotted as a function of the cadence and AMPO. For all conditions, it was found that AMPO^−^ was substantial, with the average across all conditions being 56,4% of imposed AMPO. AMPO^−^ relative to AMPO became particularly large at high cadence and low AMPO. In line with our reasoning, AMPO^−^ increased with increasing cadence. However, contrary to our expectations, AMPO^−^ did not consistently increase with increasing AMPO.

**Figure 3.**
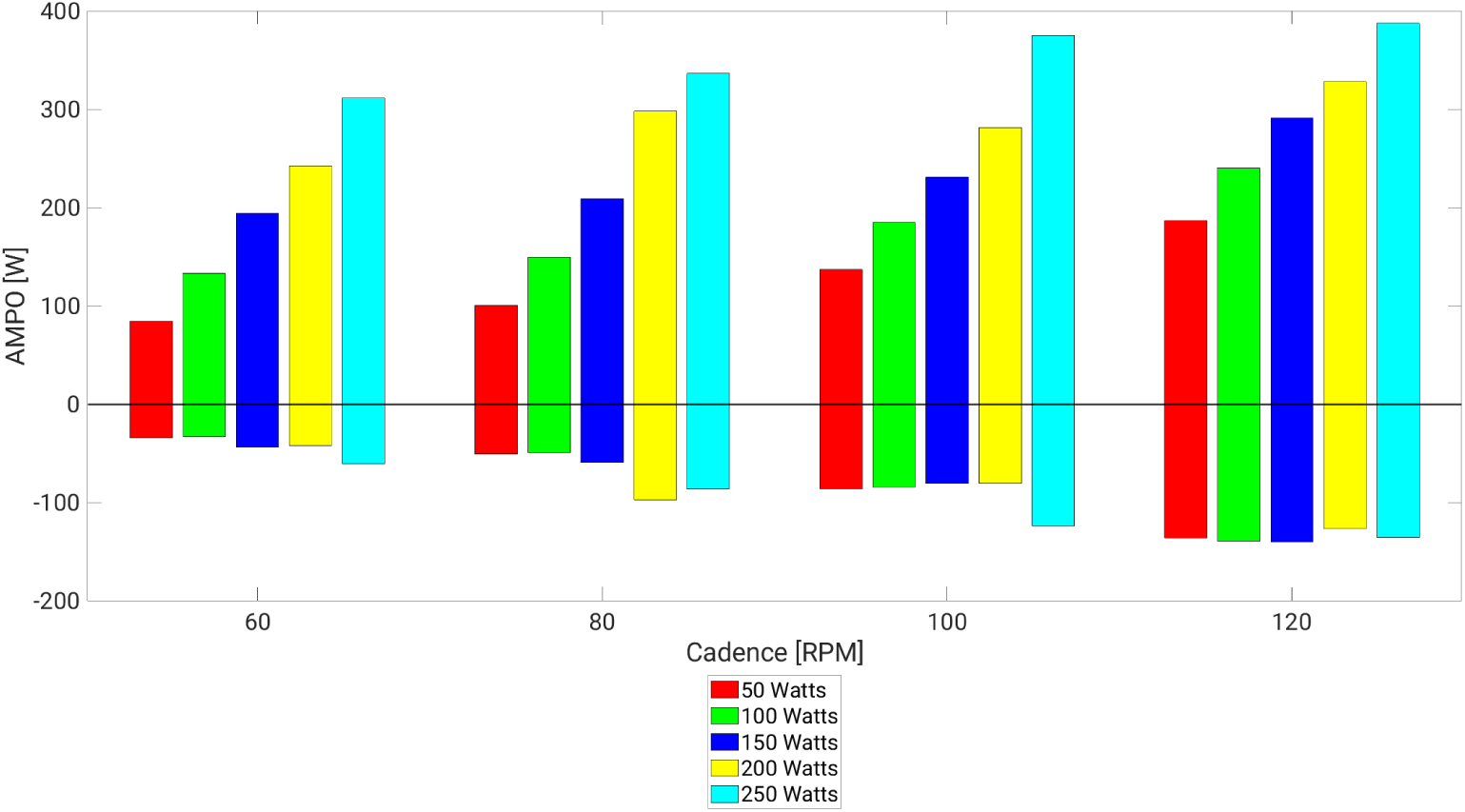
AMPO ^+^ and AMPO ^−^ as a Function of Cadence and AMPO.

Fig. 4 shows the muscle forces as a function of the crank angle at a cadence of 80RPM. In line with van Soest and Casius (2000), we found that negative mechanical muscular power was caused by the need for force buildup to start during the preceding lengthening phase. See, for example, the vasti at crank angles before 360^◦^ or the iliopsoas before 180^◦^ in Fig. 4.

**Figure 4.**
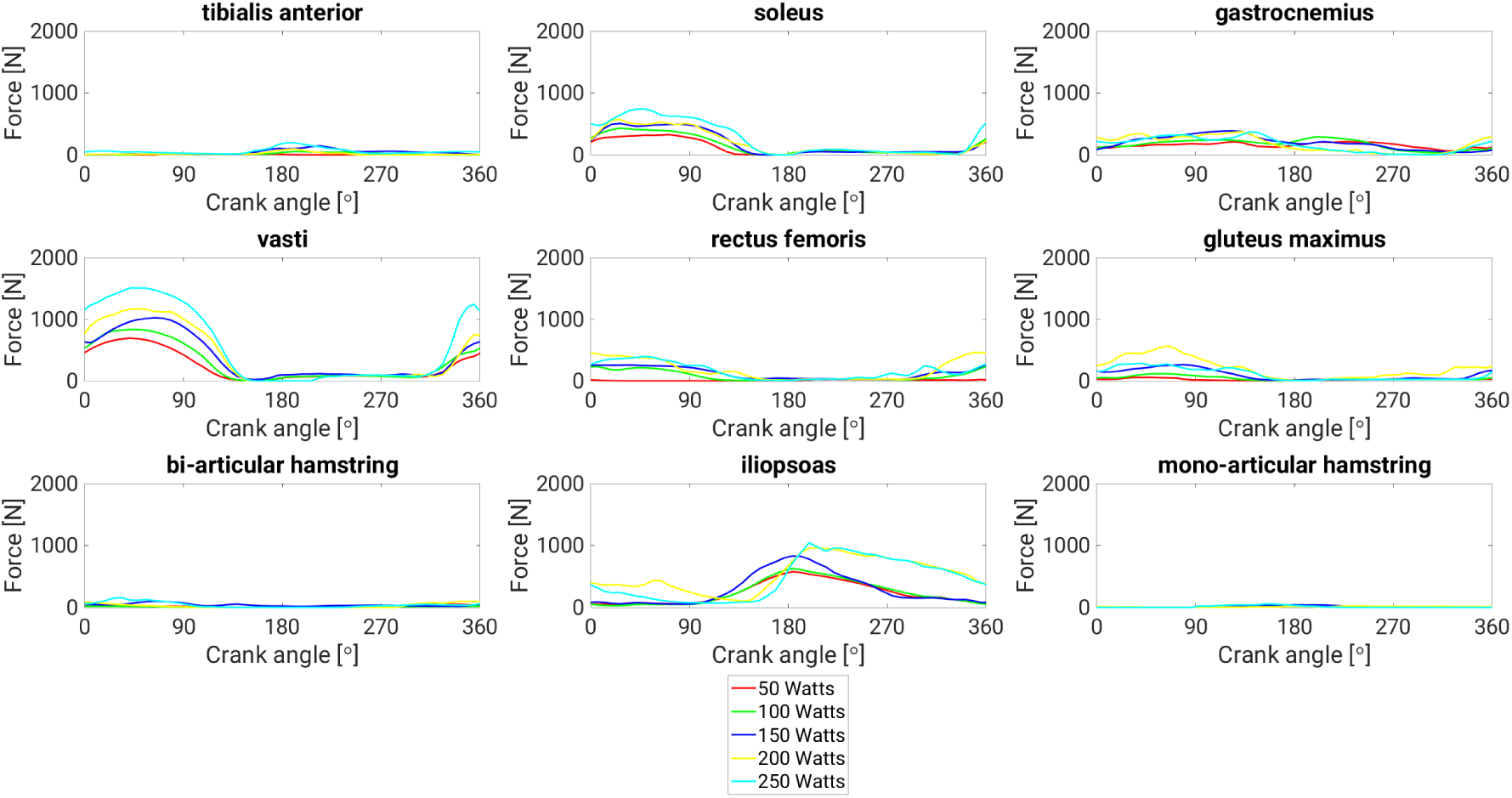
Muscle Force as a Function of Crank Angle at Different AMPO Levels. Force of the contractile element as a function of the crank angle (0*^◦^* is top-dead center) at a cadence of 80RPM.

### 3.3 AMPO*^−^* had a large effect on GE

Fig. 5 shows GE values reported in the literature (Chavarren & Calbet, 1999; Gaesser & Brooks, 1975; Samozino et al., 2006; Sidossis et al., 1992) before and after correcting for the effects of AMPO^−^. The corrections led to a substantial increase in GE, with average GE being 16.9% before and 27.5% after.

**Figure 5.**
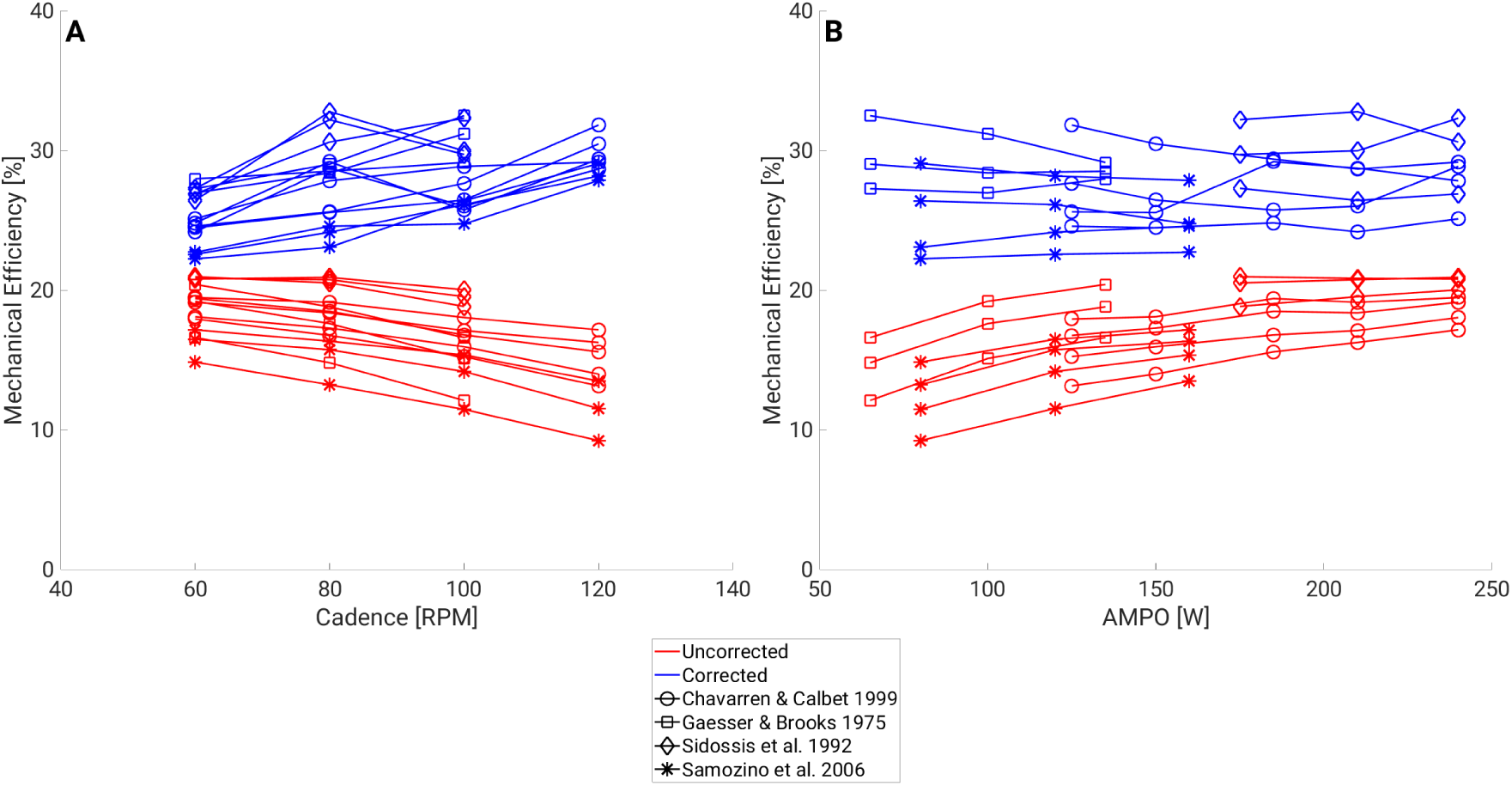
GE With and Without Correcting for AMPO ^−^. GE during sub-maximal cycling with and without correcting for the effects of AMPO*^−^*, plotted as a function of either the cadence (**A**) or AMPO (**B**). Represented studies are Chavarren and Calbet (1999), Gaesser and Brooks (1975), Samozino et al. (2006), and Sidossis et al. (1992).

In Fig. 5, it can also be seen that the relationships between cadence, AMPO and GE completely changed after correcting for AMPO^−^. Before the application of the corrections, GE consistently decreased with increasing cadence, and increased with increasing AMPO. After correcting for negative mechanical muscular power, both dependencies of GE disappeared.

## 4 Discussion

### 4.1 Overview

In this study we showed that AMPO^−^ can be substantial during sub-maximal cycling with the average across conditions being 56,4% of imposed AMPO. This is an important finding, as current estimates of GE measured in sub-maximal cycling typically assume that muscles do not produce negative mechanical muscular power. We showed that AMPO^−^ will have a considerable influence on mechanical efficiency estimates made during sub-maximal cycling. Correcting GE increased the average from 16.9% to 27.5%. Furthermore, we showed that corrected GE did not depend on cadence or AMPO. In other words, the experimentally observed relationship between GE, cadence and AMPO could potentially be explained by negative mechanical muscular power. Before discussing our findings, we will first investigate the sensitivity of our results with respect to the cost function. Then we will discuss the relationships between cadence, AMPO and GE in light of our findings. Lastly, we will describe the implications of our results in regards to current estimates of the *in vivo* muscular mechanical efficiency.

### 4.2 The influence of the cost function on the cycling pattern

To test the sensitivity of our results with respect to the cost function, we repeated our optimizations with a different cost function based on AMPO^+^. The results were then compared with those from the fatigue cost function. AMPO^+^ was chosen for the cost function, because minimization of AMPO^+^ given an imposed AMPO will result in the minimization of AMPO^−^ as well. The cost function was defined as follows:

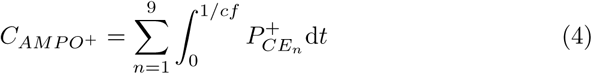

Where 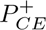 was the instantaneous positive mechanical power of the contractile element, with 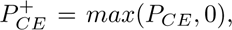 thus equalling *P_CE_* when it was positive, and otherwise being equal to 0. The max-function has a discontinuous derivative, which is undesirable when solving numerical optimization problems. We replaced the max-function with a smooth approximation:

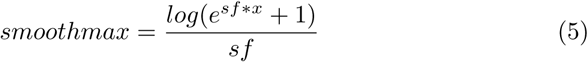

Where *sf* was a smoothness factor equalling 1.

As can be seen in Fig. 6, AMPO^−^ was substantially smaller when using the AMPO^+^ cost function. To explain high AMPO^−^ when using the fatigue cost function, we look at STIM as a function of crank angle (see Fig. 7). In general, minimizing fatigue strongly penalizes high STIM. This results in an optimal STIM pattern that tends to be low, but not zero, for prolonged durations, which leads to the production of negative mechanical muscular power.

**Figure 6.**
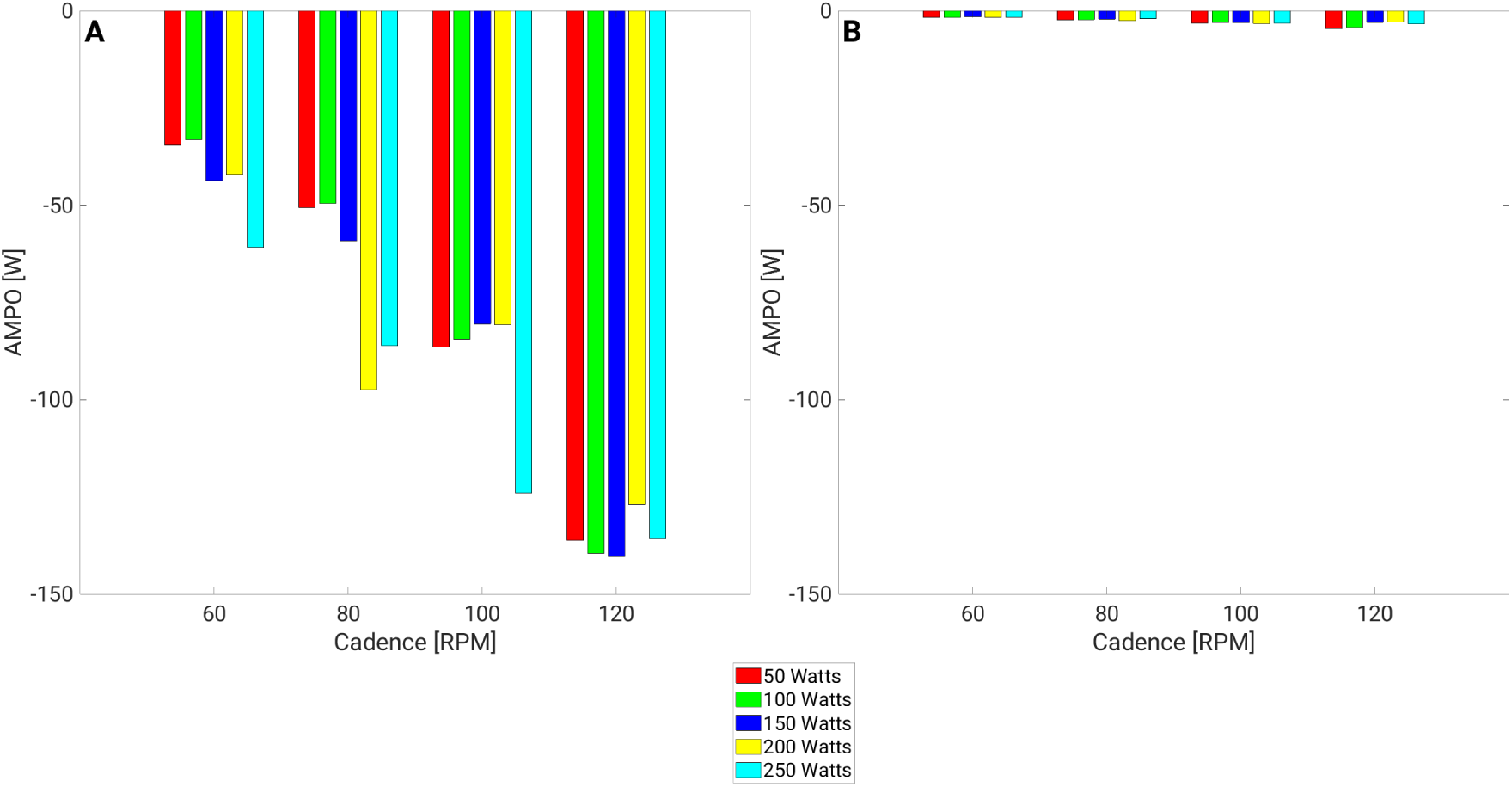
AMPO ^−^ Using Muscle Fatigue or AMPO^+^ as the Cost Function. AMPO*^−^* as a function of the cadence and AMPO, using either muscle fatigue (**A**) or AMPO^+^ (**B**) for the cost function.

**Figure 7.**
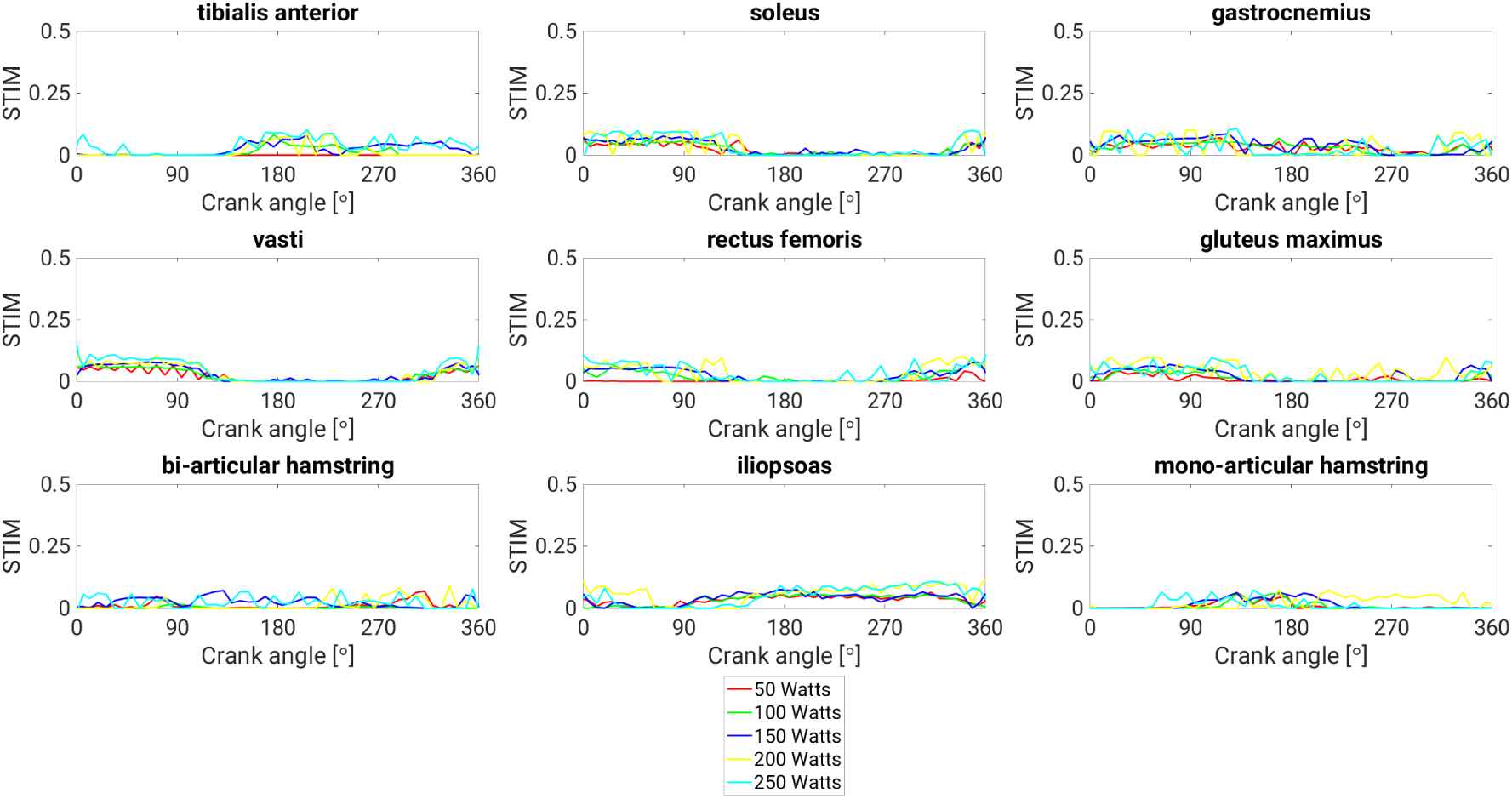
STIM as a Function of Crank Angle at Different AMPO Levels. STIM as a function of the crank angle (0*^◦^* is top-dead center) at a cadence of 80RPM.

Fig. 8 and 9 show illustrations of the kinematics obtained with the fatigue and AMPO^+^ cost functions. These figures show that the ankle was kept in a plantar flexed position when using the AMPO^+^ cost function. Comparing Fig. 10 and 11 we can see that only 2 muscles, the gluteus maximus and iliopsoas, produced substantial mechanical power when using the AMPO^+^ cost function. Compared to the fatigue cost function, the kinematics and mechanical muscular power patterns obtained with the AMPO^+^ cost function match poorly with experimental data (Gidley et al., 2019; Jorge & Hull, 1986; Pouliquen et al., 2021; Wozniak Timmer, 1991; Yum et al., 2021). This leads to an interesting point: if humans indeed try to minimize musle fatigue, as suggested by Gidley et al. (2019), then they are likely to produce a substantial amount of negative mechanical muscular power. Here, we would like to note that other cost functions related to muscle stimulation, e.g. control effort (*ST IM* ^2^, see Diedrichsen et al. (2010)), or to muscle force 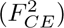 yield similar results in terms of AMPO^−^ (results not shown).

**Figure 8.**
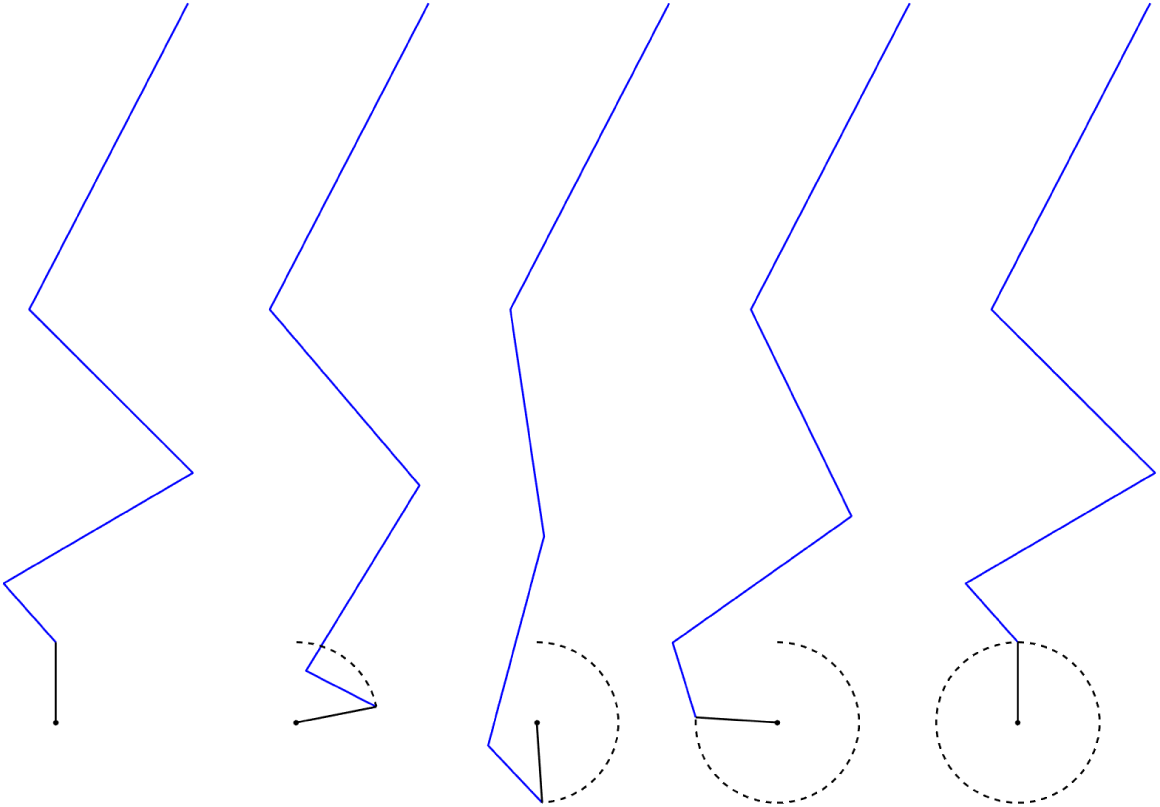
Illustration of a Cycling Pattern Obtained Using Muscle Fatigue as the Cost Function.

**Figure 9.**
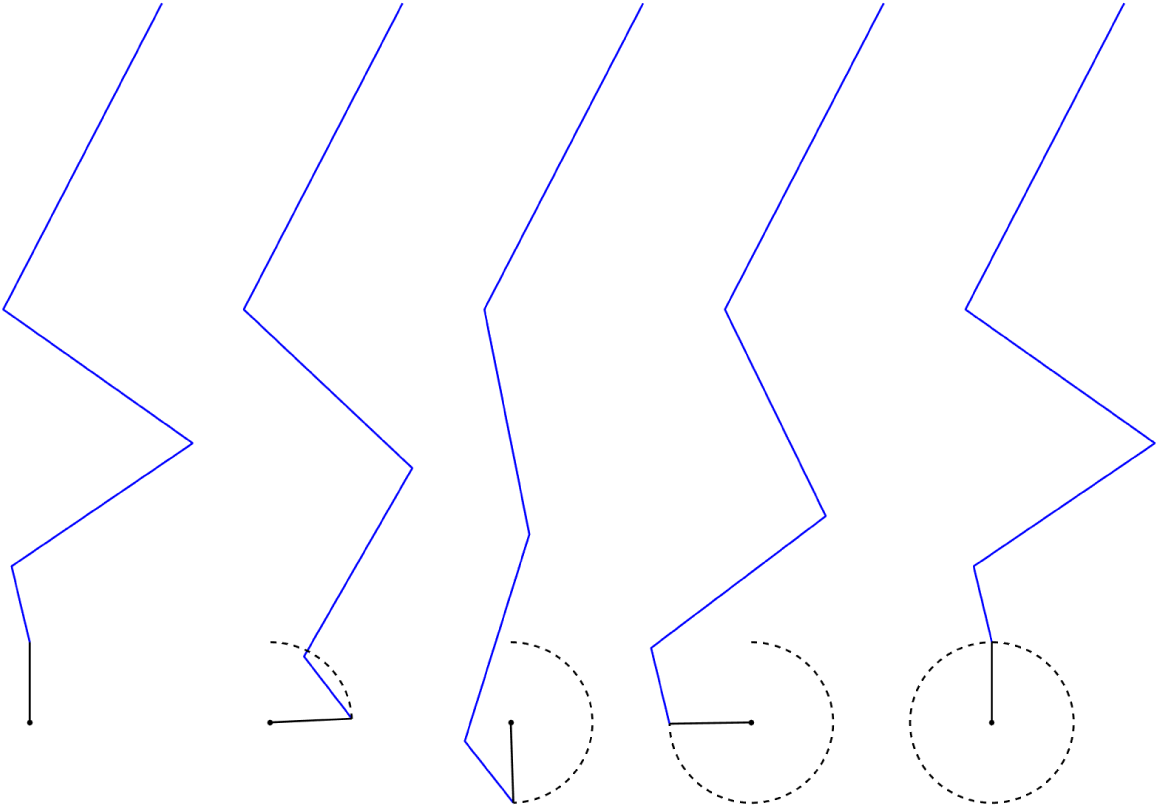
Illustration of a Cycling Pattern Obtained Using AMPO ^+^ as the Cost Function.

**Figure 10.**
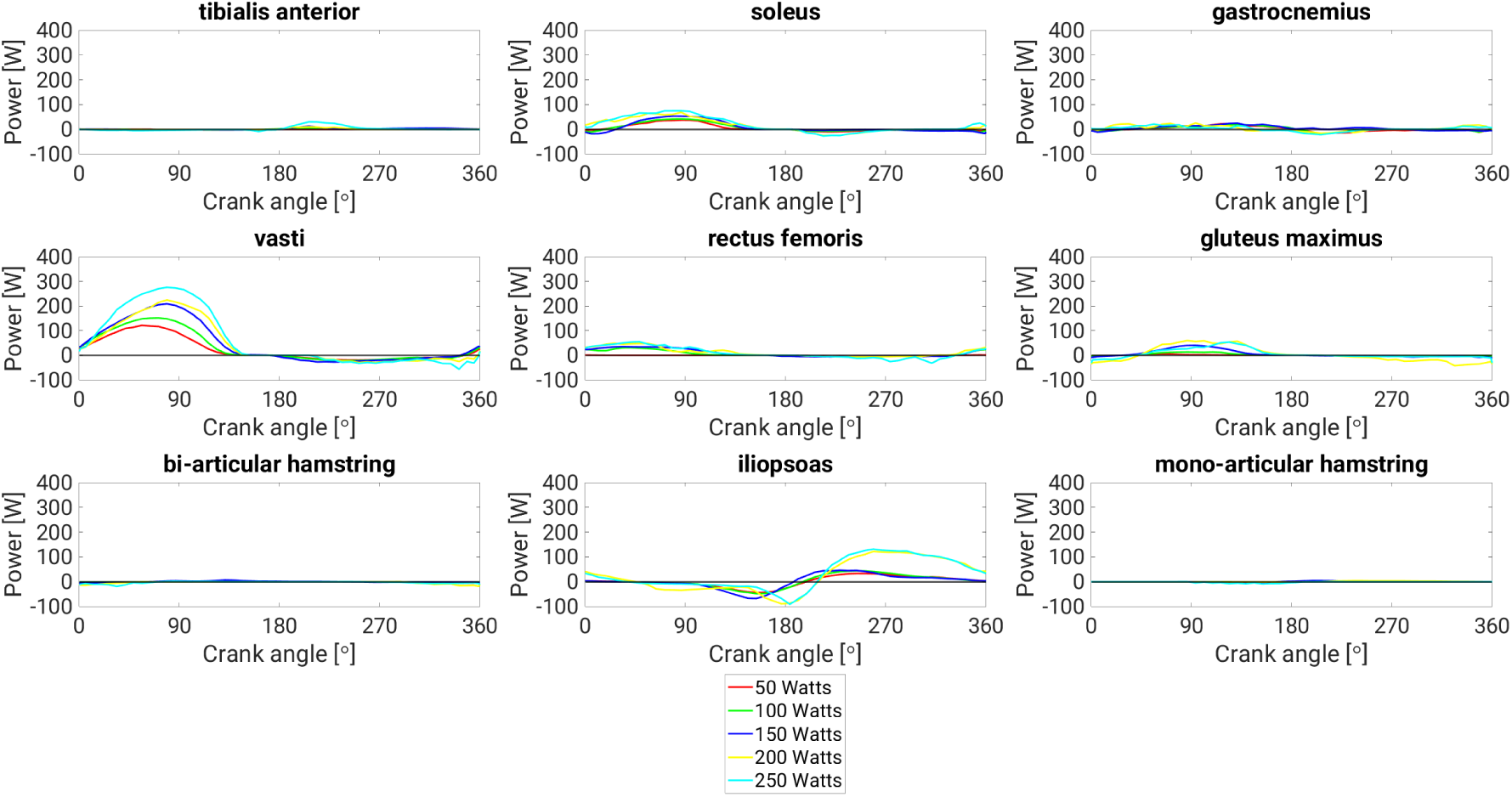
Muscular Mechanical Power as a Function of Crank Angle Using Muscle Fatigue as the Cost Function. Mechanical power of the contractile element as a function of the crank angle (0*^◦^* is top-dead center) at a cadence of 80RPM.

**Figure 11.**
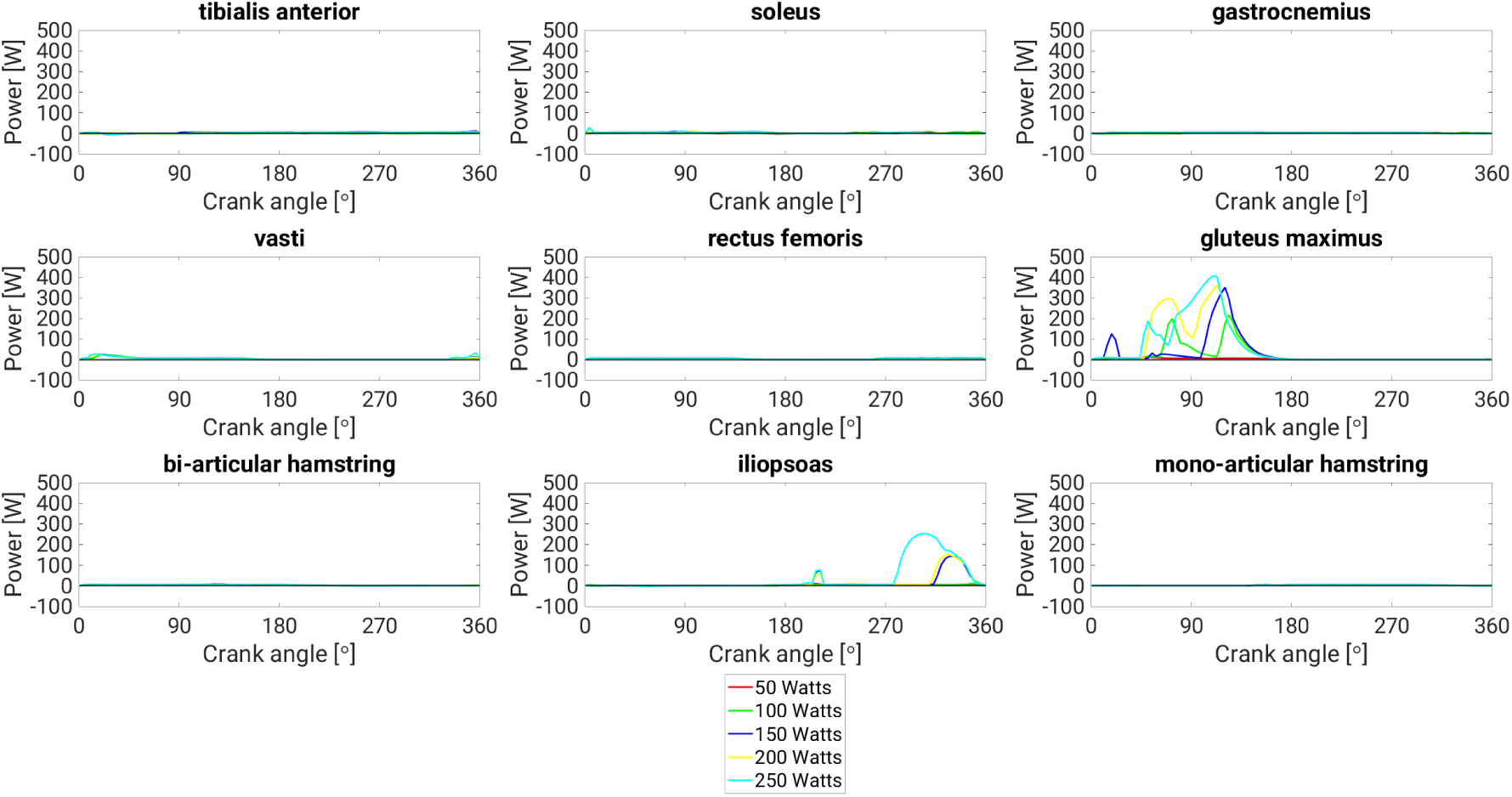
Muscular Mechanical Power as a Function of Crank Angle Using AMPO ^+^ as the Cost Function. Mechanical power of the contractile element as a function of the crank angle (0*^◦^* is top-dead center) at a cadence of 80RPM.

### 4.3 Re-examining the effects of cadence and AMPO on GE

Previous studies have reported a decrease in experimentally observed GE with increasing cadence (Chavarren & Calbet, 1999; Gaesser & Brooks, 1975; Sallet et al., 2006; Samozino et al., 2006). As explained in the Introduction, this finding is in line with our expectation that AMPO^−^ would increase with increasing cadence. As shown in Fig. 3, AMPO^−^ did indeed increase with increasing cadence. When correcting GE for AMPO^−^ (and thus for AMPO^+^ given an imposed net AMPO; see Methods), we found that the relation between GE and cadence disappeared (see Fig 5).

It was expected that an increase in AMPO would result in an increase in AMPO^−^. However, as shown in Fig. 3, we did not find a consistent effect of AMPO on AMPO^−^. As can be seen in Fig. 7, STIM(*t*) patterns were very similar across AMPO levels. Similar STIM(*t*) levels during the lengthening phase across AMPO levels could therefore lead to similar amounts of AMPO^−^. Interestingly, a relatively constant AMPO^−^ across AMPO levels is in line with the experimental finding that GE increases with increasing AMPO. The same absolute amount of AMPO^−^ is relatively smaller at higher AMPO levels, and thus has a smaller effect on GE, allowing for higher GE at higher AMPO levels. Again, when correcting for AMPO^−^, the experimentally observed relationship between GE and AMPO disappeared (see Fig. 5).

### 4.4 Implications for measures of the *in vivo* muscular mechanical efficiency in cycling

It is important to point out that our findings regarding the effects of AMPO^−^ are not limited to GE. Much of the discussion about the usage of different measures of mechanical efficiency that use AMPO, such as the net efficiency, work efficiency or delta efficiency, has focused on the used estimate of the muscular metabolic power, and more specifically on the validity of the used base-line subtraction (Ettema & Lorås, 2009; Stainbsy et al., 1980). However, all of these measures use AMPO for their estimates of AMPO^+^. Our results suggest that even if one of these measures were to use an appropriate estimate of the muscular metabolic power, then the muscular mechanical efficiency would still be underestimated due to errors in the estimated AMPO^+^. As far as we are aware, no previous studies have questioned the validity of the usage of AMPO as an estimate of AMPO^+^ in sub-maximal cycling. We also did not find any studies in which *in vivo* estimates of the muscular mechanical efficiency were corrected for the influence of AMPO^−^.

To summarize, we believe that muscular mechanical efficiency during submaximal cycling is likely higher than previously thought, due to the neglect of the influence of AMPO^−^ on mechanical efficiency estimates in previous studies. Furthermore, muscular mechanical efficiency likely depends less on the cadence and AMPO than has previously been reported. Part of the variations in mechanical efficiency estimates due to cadence and AMPO can be explained by the dependency of AMPO^−^ on the cadence and AMPO.

## 5 Conclusion

Using a simulation model of a human cyclist, we have estimated AMPO^−^ during sub-maximal cycling. We found that AMPO^−^ was considerable in all conditions. Correcting experimentally observed GE values reported in the literature for the effects of AMPO^−^ led to substantial increases in GE values. These findings suggest that current measures used to estimate the *in vivo* mechanical efficiency of muscle during sub-maximal cycling underestimate the true muscular mechanical efficiency.

